# Single-cell analysis of chromatin and expression reveals age- and sex-associated alterations in the human heart

**DOI:** 10.1101/2022.07.12.496461

**Authors:** David F Read, Gregory T Booth, Riza M Daza, Dana L Jackson, Rula Green Gladden, Sanjay R Srivatsan, Brent Ewing, Jennifer M Franks, Cailyn H Spurrell, Anne Roshella Gomes, Diana O’Day, Aishwarya A Gogate, Beth K Martin, Lea Starita, Yiing Lin, Jay Shendure, Shin Lin, Cole Trapnell

## Abstract

Sex differences and age-related changes in the human heart at the tissue, cell, and molecular level have been well-documented and many may be relevant for cardiovascular disease. However, how molecular programs within individual cell types vary across individuals by age and sex remains poorly characterized. To better understand this variation, we performed single-nucleus combinatorial indexing (sci) ATAC- and RNA-Seq in human heart samples from nine donors. We identify hundreds of differentially expressed genes by age and sex. Sex dependent alterations include pathways such as TGFβ signaling and metabolic shifts by sex, evident in both transcriptional alterations and differing presence of transcription factor (TF) motifs in accessible chromatin. Age was associated with changes such as immune activation-related transcriptional and chromatin accessibility differences, as well as changes in the relative proportion of cardiomyocytes, neurons, and perivascular cells. In addition, we compare our adult-derived ATAC-Seq profiles to analogous fetal cell types to identify putative developmental-stage-specific regulatory factors. Finally, we train predictive models of cell-type-specific RNA expression levels utilizing ATAC-Seq profiles to link distal regulatory sequences to promoters, quantifying the predictive value of a simple TF-to-expression regulatory grammar and identifying cell-type-specific TFs.

## Introduction

Profound alterations in cardiac function and disease risk have long been evident at the level of individuals’ traits such as sex ^1^ and age ^2^. For example, female hearts exhibit more modest declines in cardiomyocyte numbers over time than males ^3^ and display distinct vascular elasticity properties ^4^, while aged hearts display ventricular hypertrophy, tissue stiffening, and inflammation ^2,5^. However, there is substantial uncertainty in the exact molecular and cellular hallmarks - much less causal mechanisms of these clinically evident, consequential differences. A robust understanding of those molecular processes could set the stage for personalized therapeutic intervention.

To achieve cell-type-resolved but high-throughput measurements of cardiac biology, single cell methods have been employed in numerous studies of human hearts and model organisms. In humans, these analyses profiled the diversity of cardiac cell types and subtypes ^6,7^ and generated genome-wide maps of cell-type specific regulatory programs ^8^. In model organisms, single cell approaches have not only generated atlases of healthy tissue ^9,10^ but have also been used in controlled experiments to dissect the alterations occurring in processes such as aging ^11^ and heart disease ^12–14^. Similar approaches have begun to profile clinically important contrasts in human samples directly, such as identifying a handful of transcripts that vary by age in the healthy human heart ^7^, variation in myeloid cell abundance in age ^15^, and alterations during disease in single-cell ATAC-Seq ^8^ and RNA-Seq ^15^ data. Further analyses and larger datasets of the human heart may unlock more extensive insights into how alterations in transcriptional and epigenetic states characterize variation between individuals and advance our understanding of the genomic programs regulating cells.

Chromatin regulation represents a significant element of specialized cell function within or between conditions. During development, transcription factors play variable roles over the course of cardiac development ^16,17^. Of clinical concern, individual transcription factors may play decisive roles in diseases such as cardiac fibrosis ^18^ while genetic variation may act through regulatory mechanisms to affect disease risk and individual variation ^19^. Parallel advances in quantitative models of gene expression ^20–22^ and extensive generation of epigenetic datasets in primary human hearts ^8,17,23^ bode well for the utility of further, diverse epigenetic datasets in revealing intra- and inter-state regulatory programs.

To extend knowledge of molecular cell-type-specific cardiac processes between and within individuals, we generated and analyzed matched single-cell ATAC-Seq (117,738 cells) and RNA-Seq (89,404 cells) datasets from 15 samples spanning 9 individuals. As a resource, our dataset contributes substantially to the catalog of single-cell profiles of the human heart. The number of individuals profiled combined with a hierarchical mixed model regression approach allows us to resolve age- and sex-dependent transcriptional and chromatin accessibility changes apart from confounding by donor-level variation. We find that transcriptional and regulatory programs display widespread variation by these covariates, observing both cell-type-specific and largely pan-celltype alterations. For example, sex was associated with alterations in transcriptional signatures of oxidative phosphorylation as well as differing accessibility at ATAC-Seq peaks containing motifs of TFs known to regulate metabolic rewiring. Furthermore, we employ ATAC-Seq data to identify putative life-stage specific TFs, finding indications of adult-specific activity by RFX family TFs in adult vascular endothelium and macrophages. Finally, we develop cell-type-specific gene expression models that utilize informative distal regulatory sites to account for approximately a quarter of transcriptional variation using a simple TF motif regulatory code.

## Results

### Single cell ATAC- and RNA-Seq library generation and cell annotation

We collected heart samples from nine healthy adult donors (**Supplementary Table 1**) with hearts collected on hemodynamic support to eliminate warm ischemic time (see “Methods” section). Samples represented four anatomical sites from the heart, though most were collected from the heart apex or left ventricular wall. In total, we prepared 15 samples for single-cell analysis, with each sample representing a single anatomical site in a particular donor. We powdered frozen tissue and split portions into aliquots for appropriate nuclei isolation and fixation for ATAC-Seq and RNA-Seq separately (**Fig. 1A**).

**Figure 1:**
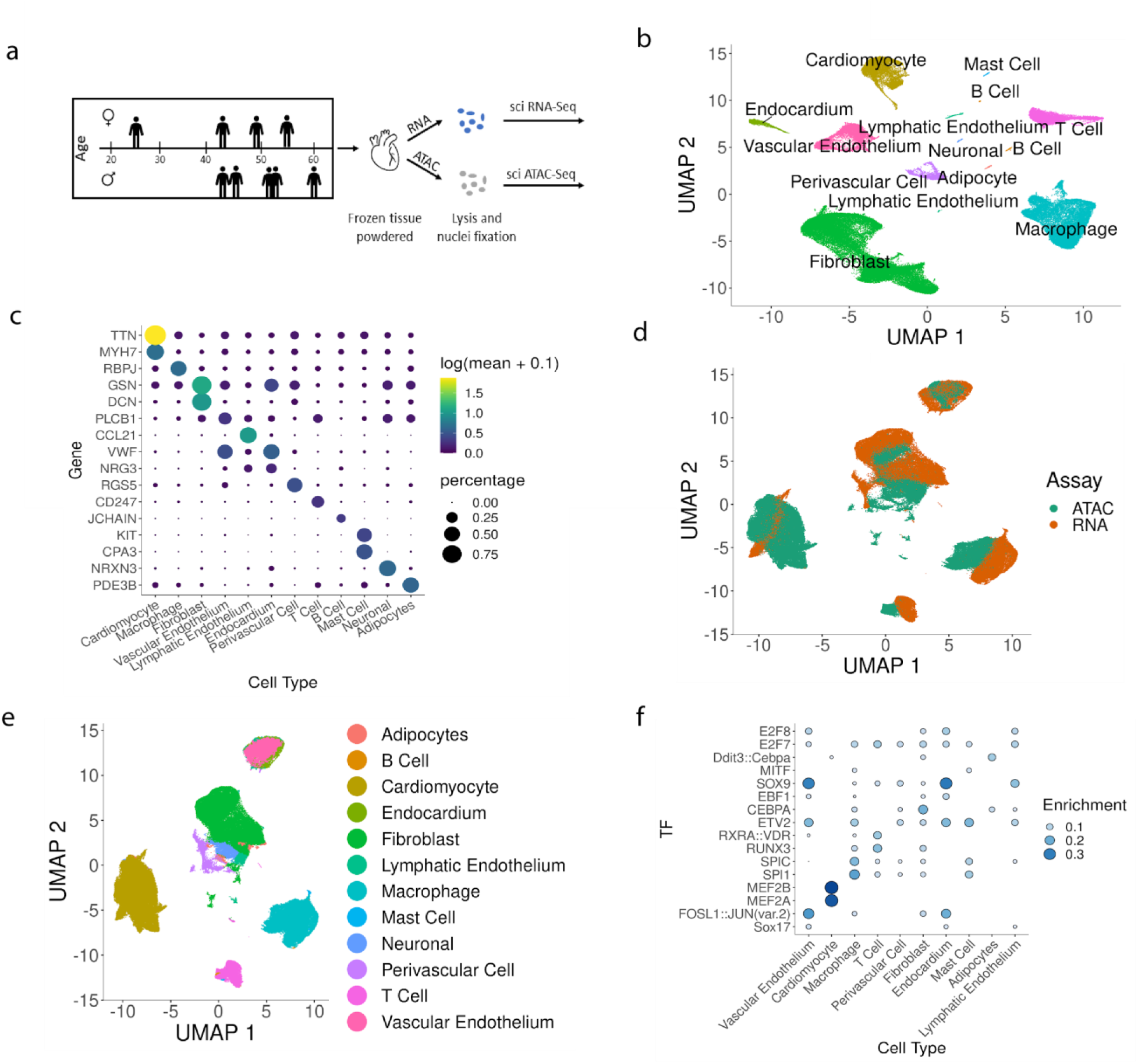
Overview of datasets. A) Experimental setup. B) UMAP of RNA-Seq data, coloring by cell type. C) Plot of marker gene expression by cell type, showing both mean expression (dot color) and the proportion of each cell type with nonzero expression for that marker (dot size). D) UMAP of a co-embedding of snRNA- and ATAC-Seq data, coloring by data type. E) UMAP of co-embedded RNA and ATAC data, coloring by the assigned cell type F) Enrichment of TF motifs in the accessible peaks of ATAC-Seq cells

We generated single-nuclei RNA-Seq libraries using 3-level sci RNA-Seq ^24^. We modified the nuclei isolation protocol to use additional RNase inhibitors, mechanical dissociation of tissue, and 5% glutaraldehyde for tissue fixation (see “Methods” section). Additionally, in order to reduce background RNA levels that commonly contribute noise to single-cell RNA-Seq data ^25^ we included a FACS sorting step following ligation. One sample failed completely (donor “W137”) but the remaining 14 samples yielded 89,404 nuclei after doublet removal and filtering. A UMAP embedding of all transcriptomes contained numerous clearly separated clusters, with contributions from distinct samples across clusters (**Supplemental Fig. S1A**). Examination of marker genes revealed that clusters corresponded to specialized cells of the human heart including cardiomyocytes, fibroblasts, macrophages, and endothelial cells (**Fig. 1B, C**). The RNA data also clearly resolves rare cell populations such as adipocytes, neuronal cells, mast cells, and B cells **(Fig. 1B**).

Separately, we prepared single-nuclei ATAC-Seq data from powdered frozen tissue using 3-level sci ATAC-Seq ^23^, generating 117,738 ATAC-Seq profiles after filtering and doublet removal. Single-cell ATAC seq data is more difficult to annotate than RNA data because open chromatin around a gene doesn’t always indicate that gene is robustly expressed ^26^. Due to this difficulty in defining cell types, we used a co-embedding approach to find a low-dimensional embedding of RNA- and ATAC-Seq data simultaneously (**Fig. 1D**), then transferred cell-type labels from RNA to ATAC profiles using a k-nearest neighbor classifier (**Fig. 1E**; see “Methods” section; see **Supplementary Fig. 2** for post-filtering QC). Using these assignments, within the ATAC-Seq data we identify strong enrichments for expected cell-type-specific transcription factors (**Fig. 1F**) such as MEF2 family transcription factors in cardiomyocytes, SPI1 (also known as PU.1) in macrophages, and CEBPA in fibroblasts, in agreement with recent analyses of adult single-cell ATAC-seq data in adult human hearts ^8^.

**Figure 2:**
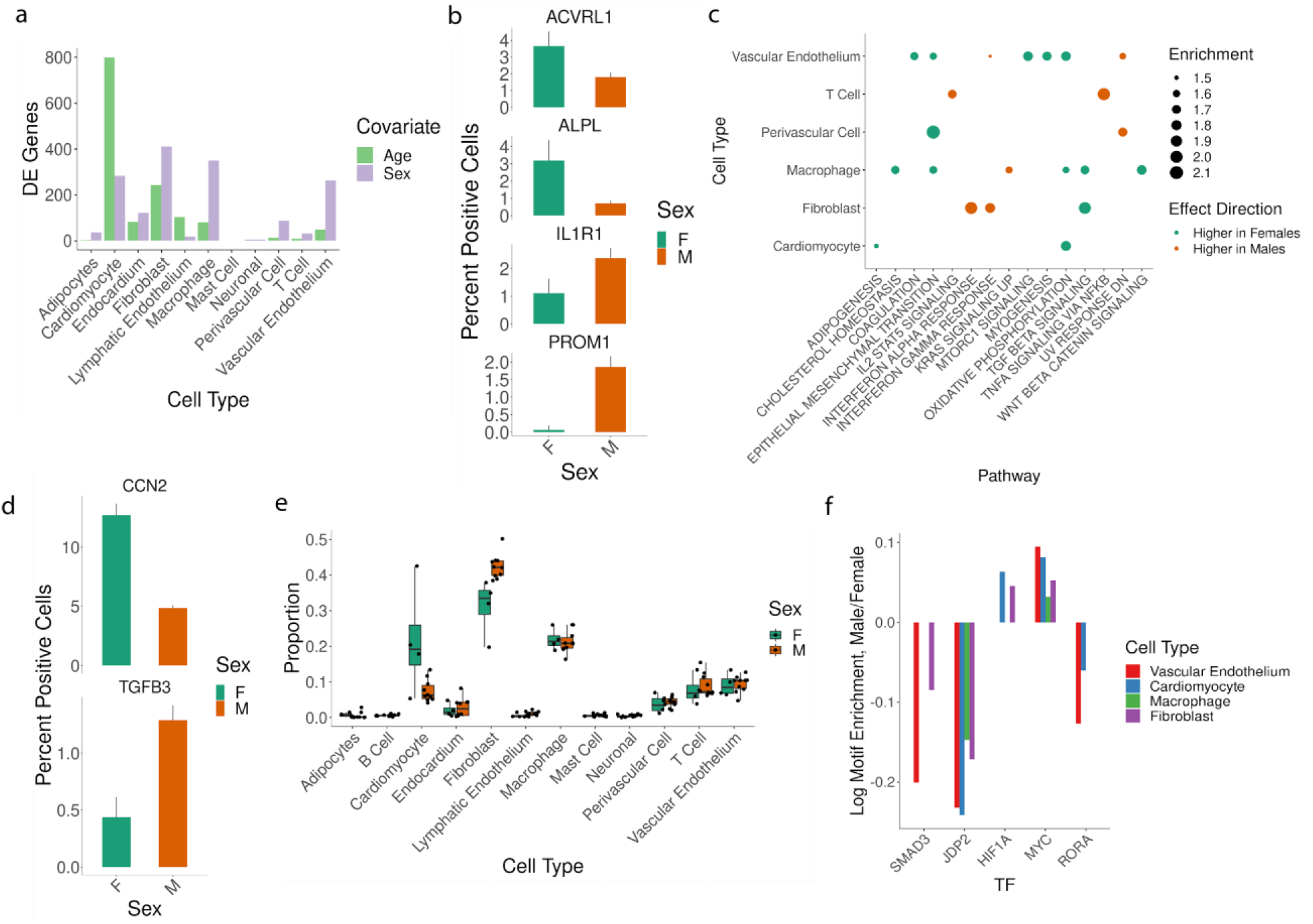
Alterations by sex in the heart. A) Total differentially expressed genes by age or sex (FDR = .1). B) Vascular endothelial cells positive for expression of four genes that were DE by sex. C) Enrichment of biological processes by sex. Pathways “Enriched in Females “ show higher-than-expected expression in female cells after controlling for other covariates, and vice-versa. D) Fibroblasts positive for expression of two genes that were DE by sex. E) The proportion of total cells per sample that were classified as each cell type. F) Enrichment for counts of TF motifs in the accessible peaks of cell types as a function of sex. Bars shown correspond to statistically significant coefficients (FDR = .1).

### Sex corresponds to cell-type-specific differences including TGF-β signaling and metabolic alterations

We explored our single cell RNA-Seq data to identify transcriptional changes associated with age or sex within individual cell types. Commonly-used single-cell differential expression methods using fixed effect regression models do not properly account for inter-sample variation and their mis-application to datasets such as ours - in which cells are not statistically independent when derived from the same donor - can dramatically inflate false discoveries ^27,28^. Consequently, we used a mixed effect modeling framework to test for differential expression (see “Methods” section). This approach allowed us to test for variation by sex and age while controlling for expression differences due to anatomical site (as a fixed effect) and donor (as a random effect). We find dozens to hundreds of differentially expressed genes by age and sex, depending on the cell type analyzed (**Fig. 2A**). Most DE genes are found in relatively abundant cell types, and we note particularly large numbers of changes by age in cardiomyocytes as well as many differences by sex in fibroblasts, macrophages, and vascular endothelium. For example, in vascular endothelial cells (**Fig. 2B**) we observe large differences between men and women in expression of *PROM1* (also known as *CD133*), a marker for endothelial progenitor cells ^29^ and proliferative vascular endothelium ^30^. As transplanted *PROM1+* cells promote vascular regeneration in a mouse model of ischemic heart injury ^29^, altered *PROM1* levels by sex may indicate variation in proliferative endothelial populations. DE genes by sex in vascular endothelium also include *IL1R*, a receptor for pro-inflammatory IL1 signaling that is a candidate therapeutic target pathway in cardiovascular diseases including acute myocardial infarction ^31^; *ALPL*, an alkaline phosphatase that promotes cardiac fibrosis ^32^; and *ACVRL1* (*ALK1*), a TGFβ superfamily co-receptor that causes vascular malformations and hemorrhage upon depletion ^33^.

To summarize high-level changes within sex- and age-specific variation, we tested for enrichment in up- and down-regulated gene sets. Statistically significant differences between male and female hearts were evident across several cell types (**Fig. 2C, Supplementary Table 2 and 3**). One recurrent alteration was a decrease in TGFB hallmarks across several cell types. We observe decreased expression of target genes of TGFB signaling in male fibroblasts and macrophages as well as decreased hallmarks of epithelial-to-mesenchymal transition - a common downstream consequence of TGFB activity ^34^ - in macrophages, vascular endothelial, and perivascular cells (**Fig. 2C**). Additionally, we find statistically significant changes in genes important in various aspects of cell metabolism, including decreased expression of cholesterol metabolism-associated transcripts in macrophages and decreased oxidative phosphorylation-related transcripts in male cardiomyocytes, vascular endothelium, and macrophages (Figure 2C). We also observe cell-type specific alterations, such as an increase in hallmarks of IL2 and TNFA signaling in male T cells (**Fig. 2C**) consistent with elevated soluble inflammatory signaling^35,36^.

We next examined whether our data contained further clues to the source of these pathway-level changes. We find statistically significant alterations in expression levels of *TGFB3* in fibroblasts (**Fig. 2D**), but counter-intuitively expression levels are higher in men than women. Apart from altered *TBFB3* in fibroblasts we do not detect significant decreases in RNA levels of *TGFB1, TGFB2*, or *TGFB3* in any other cell type, but potentially find other between-sex differences that could affect TGFB signaling. For example, in male fibroblasts we do find decreased levels of *CCN2*, a promoter of TGFB signaling ^37^.

As it is also plausible that differences in the abundance of cell types with active roles in intercell communication could underpin male/female differences, we examined the cell type proportions of samples by donor sex. Broadly, cell type proportions were consistent between the sexes. However, we observe some differences in cell proportions by sex including as a statistically signifcant increase in fibroblast proportions in male donors (**Fig. 2E**) using a beta-binomial model accounting for variation due to sex, age, and anatomical site (**Supplementary Table 4**).

In order to additionally study regulatory programs whose activity differed by donor sex, we looked for alterations in TF motifs accessibility between male and female cells in order to validate and expand upon the observed transcriptional differences (**Supplemental Table 5)**. Consistent with our observation of decreased TGFB signaling in transcriptional data, we see reduced abundance of motifs for SMAD3 - a downstream effector of canonical TGFβ signaling - in male fibroblasts, vascular endothelial cells, and macrophages (**Fig. 2F**). We further see statistically significant decreases in accessible motifs corresponding to JDP2, a transcriptional repressor tied to alteration of TGFB1 induced EMT and fibrosis ^38,39^. Consistent with decreased expression of hallmarks of oxidative phosphorylation in males (**Fig. 2F**), we detect sex-specific changes across multiple cell types in HIF1A, MYC, and RORA, TFs known to promote glycolysis over oxidative phosphorylation ^40^. These ATAC-Seq-based analyses align with the transcriptional decreases in TGFβ signaling and oxidative phosphorylation genes we observed in male cells and identify putative regulatory mediators of sex-specific distinctions.

### Immune activation increases with age through multiple pathways

We next explored cell-type specific changes in the expression of hundreds of genes that varied by donor age (**Fig. 2A**). To understand the broad changes these alterations represent, we again tested for enrichments in gene sets for age-related expression changes within individual cell types. We found a variety of alterations, including changes in several metabolic and cell-signaling pathways (**Fig. 3A**). We again observed differences in TGFB signaling genes, with increased TGFB hallmarks increasing in aged fibroblasts and epithelial-to-mesenchymal transition-associated transcripts elevated in aged fibroblasts, macrophages, endocardial cells, and cardiomyocytes (**Fig. 3A)**. Age is associated with an increase in several immune pathways across several cell types, including hallmarks of inflammation in fibroblasts and macrophages and increased interferon response in macrophages and vascular endothelium (**Fig. 3A**). Statistically significant alterations include age-dependent increases in fibroblast expression of chemerin receptor *CMKLR1* (**Fig. 3B)**, a receptor that promotes inflammatory responses ^41^, mediates macrophage retention ^42^, and is positively correlated with atherosclerotic lesions ^43^.

**Figure 3:**
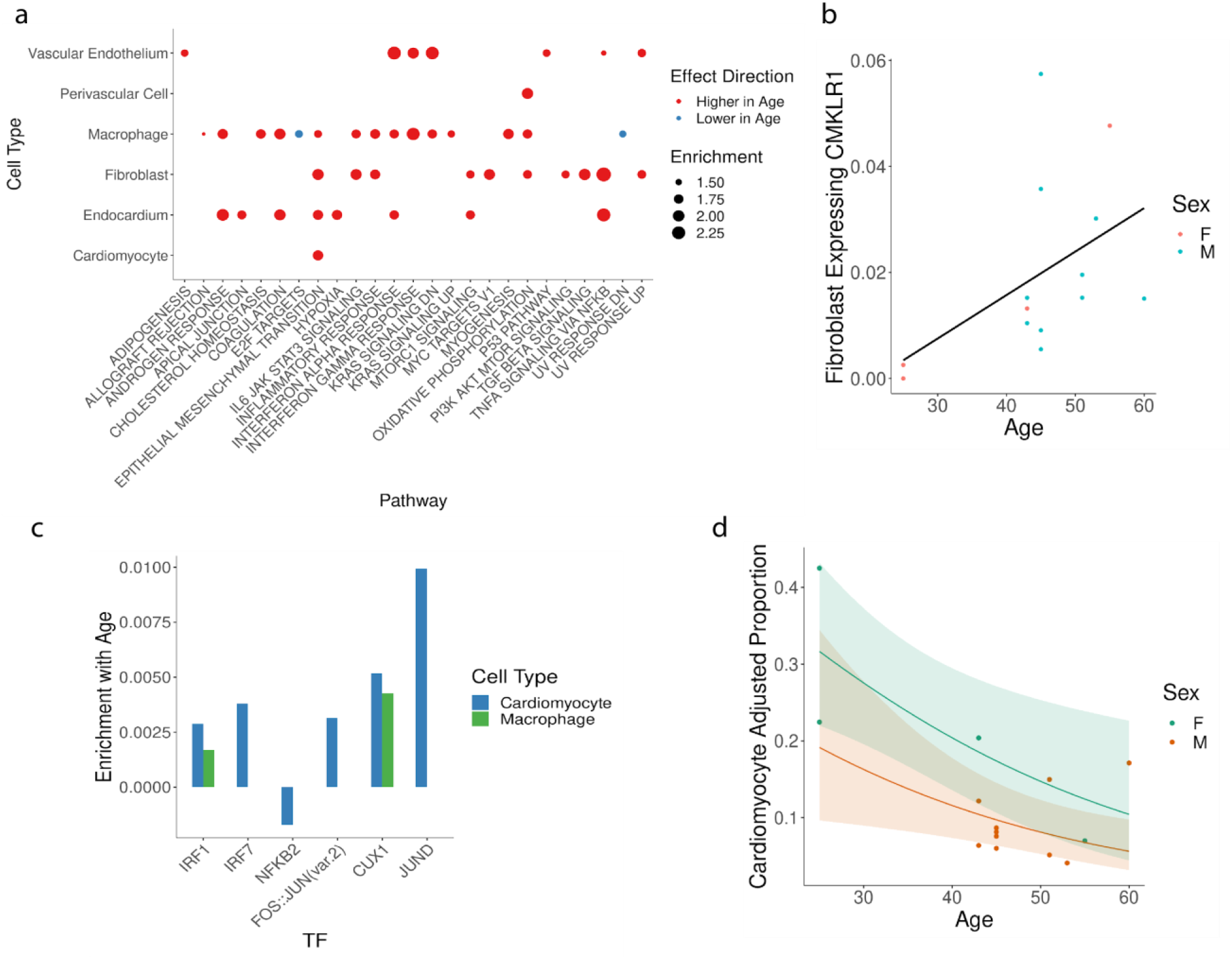
Alterations by age in the heart. A) Enrichment of biological processes by age in individual cell types. Pathways “Higher in Age “ indicates higher-than-expected expression in older cells after controlling for other covariates, and vice-versa. B) Percent of fibroblasts that are positive for CMKLR1 expression, by age of donor. C) Enrich ment for counts ofTF motifs in the accessible peaks of cell types as a function of age. Bars shown correspond to statistically significant coefficients (FDR = .1). D) Age of sample donor compared to the adjusted proportion (modified by the effect of anatomical site on proportion in a beta-binomial model) of all cells in that sample that were annotated as cardiomyocytes. Trend line shown based on predicted proportion of male (orange) and female (green) samples over a range of ages. Error range shows +/-2 standard errors.

In order to see if age-dependent alterations in immune activation hallmarks were evident at the level of chromatin remodeling, we tested for motif enrichment in accessible peaks as a function of age (**Supplementary Table S6**). We detected enrichment of IRF1 and IRF7 motifs in accessible peaks of cardiomyocytes, fibroblasts, macrophages, and vascular endothelial cells (**Fig. 3D**) consistent with increases in interferon response pathways observed in our transcriptional data (**Fig. 3A**). In addition, we observe a statistically significant decrease in accessible motifs for NFKB2 (**Fig. 3D**), a central mediator of inflammatory signaling ^44^. These changes in accessibility of motifs corresponding to key mediators of immune activation corroborate observed transcriptional changes in immune-related pathways (**Fig. 3A**).

Given heightened immune activation with age (**Fig. 3A/B)**, we wondered if increased cell senescence - a state of arrested proliferation with release of inflammatory mediators ^45^ - was evident in our samples. Senescence occurs across organs ^46^ and is suspected to play a role in age-related susceptibility to a host of cardiovascular diseases ^47^. While a clear understanding of senescence markers and causative mechanisms remains elusive ^47^, individual markers have been implicated in cardiac health. For example, the senescence marker p53 ^47^ promotes inflammation in a mouse pressure overload model ^48^ while p53 knockout reduces age-related cardiac dysfunction in mice ^49^. In all cell types we do not detect statistically significant transcriptional changes of pro-senescence regulators *P53* or *CDKN1A* (p16) ^45^. However, we find increased abundance with age in accessible peaks of cardiomyocytes and macrophages for CUX1 motifs (**Fig. 3C**), a TF linked to senescence in vasculature by promoting transcription of p16 ^50^, a cyclin dependent kinase that promotes cell cycle arrest in senescence ^51^. We additionally see increases in JUND motifs in aged cardiomyocytes (**Fig. 3C**) a factor described as a hallmark of senescence in aging ^46^. Thus, we observe cell-type specific alterations in TF motif accessibility with age for a handful of regulators associated with senescence. Future single-cell analyses - along with a deeper understanding of senescence markers - will clarify the extent and cell-type specificity of senescence in the aging human heart.

As alterations in cell type abundance - such as declines in cardiomyocyte numbers ^52,53^ and loss of cardiac stem cells ^54^ - are a characteristic of cardiac aging, we looked for alterations in cell type proportions by donor age (**Supplementary Table 4**). As expected, we observe a statistically significant decrease in the proportion of cardiomyocytes captured by donor age (**Fig. 3D**). We additionally observe alterations in other cell types, such as increases in the proportion of neuronal and perivascular cells with age (**Supplemental Fig. 3A,B**). Our observations suggest that changes in cellular proportions over time may be widespread. As cardiac function entails complex inter-cell interactions ^55,56^, such alterations may manifest in significant functional changes through disrupting intercellular signaling in addition to altering functions carried out by an individual type.

### Contrasting TF motif enrichments identify putative adult- and fetal-specific regulators

To study the global relationship between TF activity in adult cells versus their embryonic counterparts, we used a regression approach to identify TF motifs enriched in accessible chromatin of specific cell types (see “Methods” section) and compared the enriched motifs in adult cell types against corresponding fetal cell types ^23^. Motif accessibility in most fetal cell types was largely maintained in the corresponding adult types (**Supplemental Table 7**), though concordance was weaker for less abundant cell types such as adipocytes, or cell types for which our matching approach was less confident (e.g. fetal perivascular cells and adult smooth muscle cells) (**Supplemental Fig. 4**).

**Figure 4:**
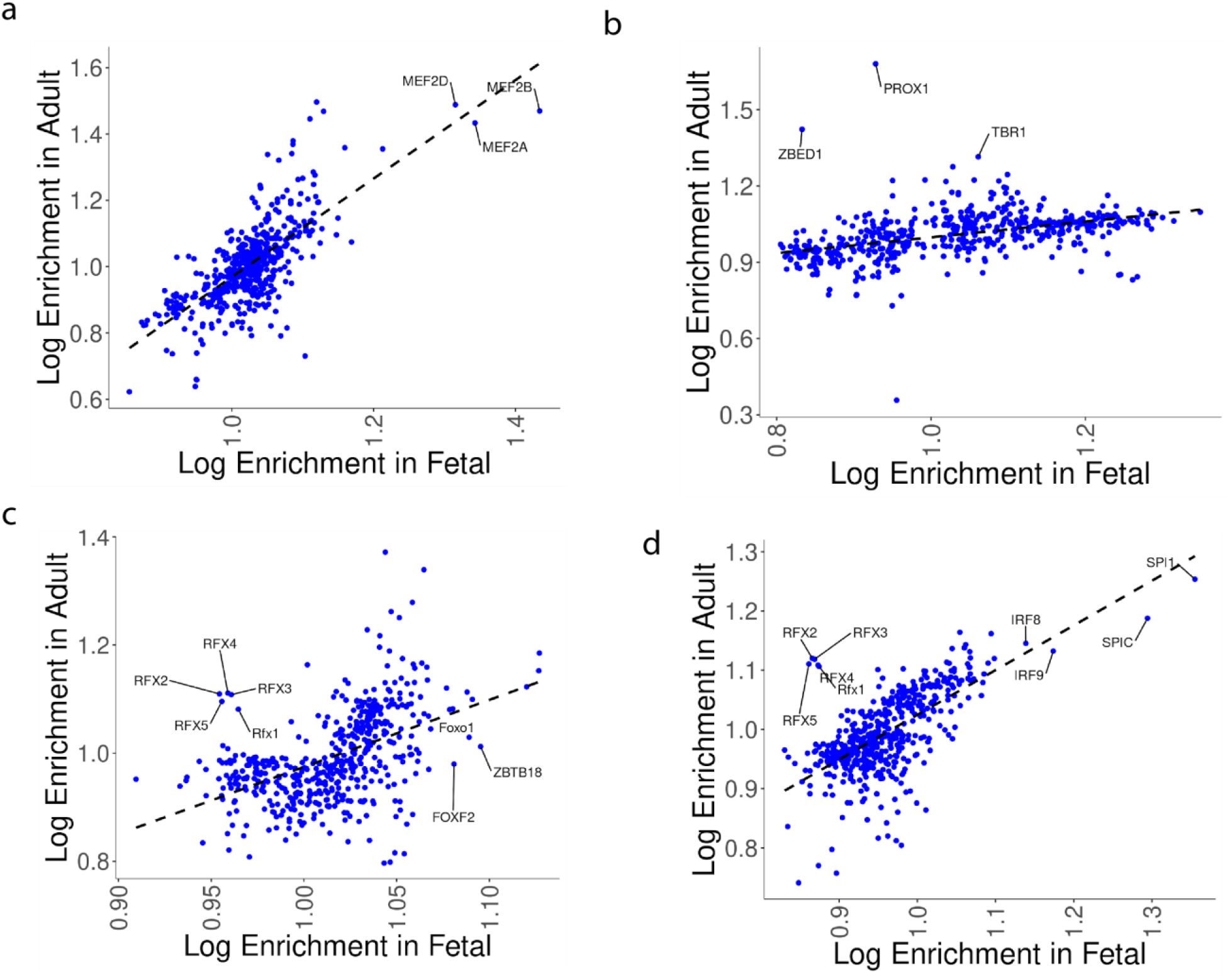
Enrichment of TF motifs in the accessible peaks of fetal or adult sn ATAC-Seq. A) Enrichments in cardiomyocytes B) Enrichments in cardiac neurons. C) Enrichments in vascular endothelial cells. Enrichments are shown for TFs that were statistically enriched in one or both of fetal or adult analyses (FDR = .1). D) Enrichments In macrophages (adult) versus myeloid lineage cells (fetal).

We next sought adult- or fetal-specific regulatory factors by looking for outliers whose enrichment was markedly higher in one developmental context versus the other. For some cell types, we see few if any obvious discrepancies in TF enrichments between fetal and adult cells. For example, TF enrichments in cardiomyocytes are highly correlated between our analysis and corresponding results from fetal data including MEF2 regulators being exceptionally enriched in accessible fetal and adult cardiomyocyte chromatin (**Fig. 4A**). This is consistent with MEF2 TFs playing a crucial role in both cardiomyocyte differentiation and maintenance ^57^.

Although the overwhelming majority of TF motifs were similarly enriched in both fetal and adult cells, apparent differences in motif enrichments occur between fetal and adult cell types. For example, while TF enrichment magnitudes are broadly correlated between adult heart and fetal neurons (Pearson correlation = .42, p =3.6e-25. **Fig. 4B**), a handful of factors show higher enrichment in adult neurons. Cardiac neurons play a pivotal role in regulating cardiac electrical and mechanical activity through a combination of intrinsic and central-nervous-system interfacing interactions ^58^, while cardiac neuron dysfunction is central in cardiac arrhythmias ^59,60^. In adult cardiac neurons we see notable enrichment for PROX1, ZBED1, and TBR1 motifs in accessible chromatin, contrasting with minimal enrichment (or depletion) of those motifs in fetal cardiac neurons (**Fig. 4B**). PROX1 plays a role in cell cycle exit and terminal differentiation of neurons in the central nervous system ^61^ while TBR1 is essential for neural specification in the developing cortex ^62^. ZBED1 plays roles in suppressing cell division ^63^, but apart from possible interactions between a ZBED1 homologue and a regulator of optic lobe formation in *Drosophila* ^*63*^, ZBED1 has not been previously characterized as a neural regulatory factor. In contrast to adult-specific motif enrichments, we observe fetal-specific enrichments of factors such as NOTO, a regulator of notochord lineage commitment ^64^, and RORA, a regulator of CNS development ^65^. Altogether, our results show that while some factors are shared between fetal and adult cardiac neurons, others may be developmentally specific.

In fetal vascular endothelial cells, we see enrichment specifically in fetal cells for known vasculature regulators FOXO1 and FOXF, both of which cause severe vascular remodeling defects and embryonic lethality upon knockout in mice ^66^ (**Fig. 4C**). In addition, we see a similar level of enrichment in accessible fetal chromatin for ZBTB18 in contrast to minimal enrichment in adult vascular endothelium (**Fig. 4C**). In the opposite direction, we see adult-specific enrichment for five RFX factor motifs (**Fig. 4C**) in adult vascular endothelium. Interestingly, we see adult-specific enrichment for these motifs in a comparison of adult versus fetal macrophages as well (**Fig. 4D**). Both of these cells types play crucial roles in vascular dysfunction ^67^, while RFX factors are correlated with epigenetic changes in hypertension patients ^68^ and RFX1 indirectly reduces monocyte recruitment in atherosclerosis ^69^. Given motif similarities between RFX factors, further work will be particularly important to understand the role of particular RFX TFs in endothelial and macrophage function. For now, our work raises the potential for adult-specific roles for RFX factors in cardiac endothelium and macrophages joining RFX factors’ previously characterized pleiotropic roles ^70^.

Overall, our comparison of cell-type specific fetal and adult motif enrichments tell us two things. First, we reproduce the observed correlation across various tissues between corresponding adult and fetal cell types’ chromatin in terms of accessible motifs ^17^. Second, motifs that are not correlated between fetal and adult cells identify candidates for developmental stage-specific regulators in cardiac cell types.

### ATAC-Seq links distal sites that improve predictive models of RNA Expression

Characterizing the regulatory roles of noncoding DNA sequences is a pressing challenge in human genetics ^71^. Although a handful of distal elements with significant functional roles in the heart have been characterized ^18^ and genome-wide maps of cis-regulatory elements have recently been published ^8^, we lack a genome-scale quantitative model of how noncoding sequences drives gene regulation. One approach to linking sequence to transcription has been to train computational models that predict each gene’s expression based on nearby sequences and/or epigenetic features ^72–76^. For example, we previously predicted gene expression based on sequence motifs in the accessible chromatin of differentiating myoblasts and found that simple transcription factor motif presence/absence explained ∼37% of transcriptional changes during differentiation ^77^. Strikingly, information from distal DNA sequences dramatically improved accuracy compared to a model that used only the promoter sequence, suggesting that much of the information needed to encode the cell-state specific expression resides in distal sequences ^77^. However, the extent to which such models generalize beyond simple *in vitro* systems to multiple *in vivo* human cell types is not clear.

To assess the potential of each cell type’s accessible chromatin to predict its transcriptome, we modeled cell-type-specific average gene expression based on promoter sequence alone or in combination with distal sites linked by ATAC-Seq information, as in our previous work ^77^. We defined hyperparameters for these cell-type specific expression models using a training set, holding aside two separate validation and test sets to measure model performance. Protein coding genes were split into train/validation/test sets at the level of whole chromosomes (see methods) in proportions of approximately 80% train/10% validation/10% test. We found that in models using only promoter sequence the best average predictive accuracy occurred when the promoter region covered 2000 bases upstream and 1000 bases downstream of a TSS (**Fig. 5A**), while use of larger or smaller regions led to inferior accuracy. After finalizing all hyperparameters (**Supplemental Fig. 5A**), two models were fit for each cell type: One used motifs absence/presence in a promoter region only as input features, while a second model used motifs found in promoters or distal DNA sites. In every cell type, models fit using proximal and distal sequence outperformed the corresponding model using promoter motifs alone, for several cell types by nearly 2-fold (**Fig. 5B**). Notably, this effect does not appear to be due simply to adding additional arbitrary sequence as use of an even larger promoter region reduced model accuracy (**Fig. 5A**).

**Figure 5:**
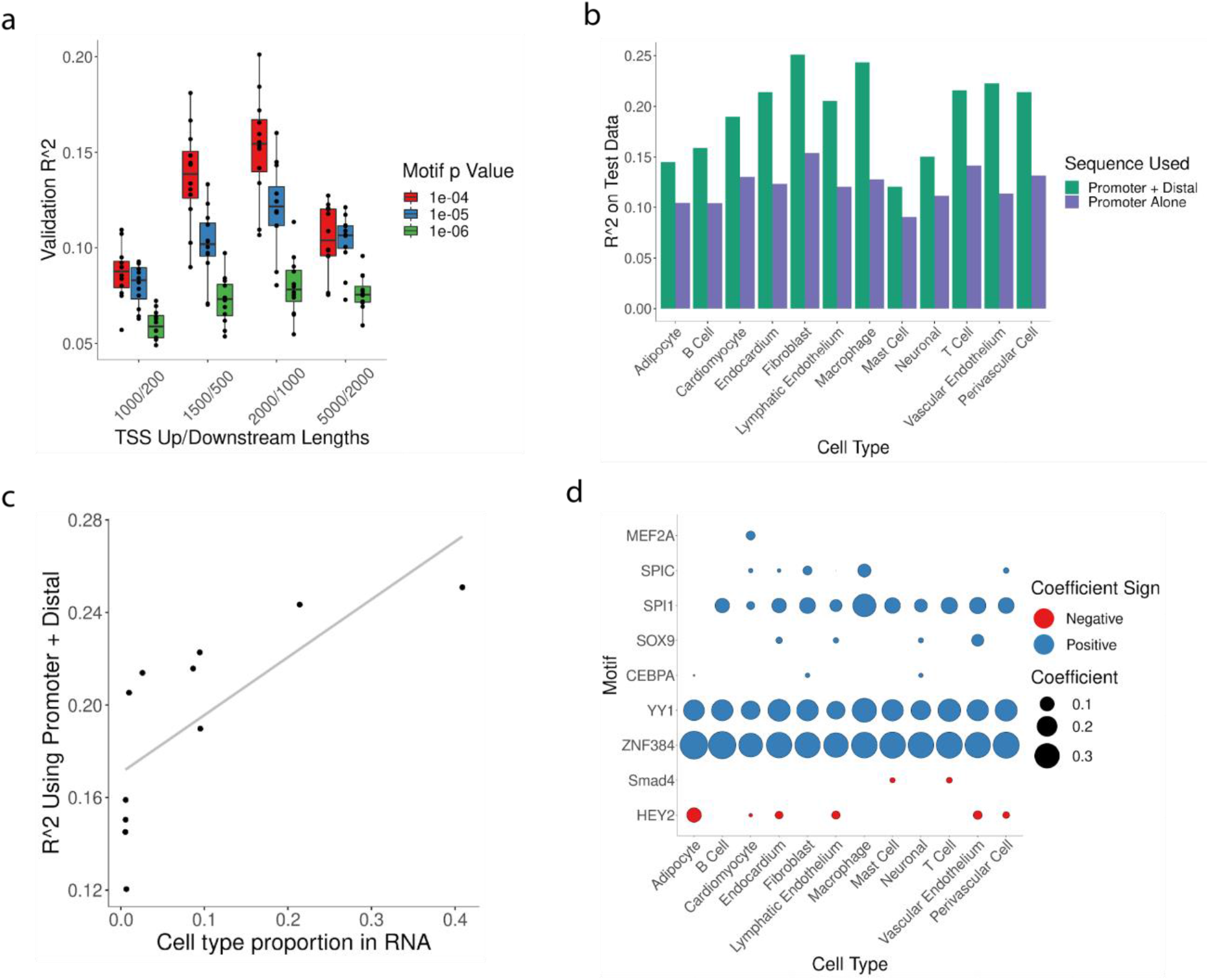
Predictive models of RNA expression. A) R^2^ for cell type-specific models (one point = one cell type) using varied p value cutoffs for calling motif presence in FIMO (color) and upstream/downstream regions (in bp) with respect to gene TSS. 8) Test set R^2^ for different cell types. R^2^ is listed for models that were trained using motifs found near gene TSS (in purple) or in either TSS or linked distal sites (in green). C) Model accuracy on test data for models using promoter and distal motifs (x-axis) and the proportion of that cell type in the sn RNA-Seq data (y-axis). D) Magnitude and direction of motif coefficients in final models fit using distal and proximal sequence.

Because models for some cell types were more accurate than others (**Fig. 5B**), we investigated if model performance was related to the abundance of the cell types.We found that cell type abundance - as quantified by a type’s proportion of total cells in RNA-Seq data - was related to accuracy for each respective cell type for models trained on promoter sequence alone (**Supplemental Fig. 5B**) or distal sequence plus promoter sequence (**Fig. 5C**). Additionally, models for abundant cell types were markedly improved by including distal information, whereas models for less abundant cell types benefitted less (**Supplemental Fig. 5C**) which suggests that collecting further data would improve our models of gene expression. For abundant cell types, a simple model that predicted each gene’s expression based on whether or not each motif was present in the accessible chromatin nearby was able to account for ∼20-25% of expression variation at the level of pseudo-bulked transcriptomes, and that around a third of that amount was due to the inclusion of distal motif information. These results demonstrate that for all cell types of the human heart for which we trained models, distal noncoding DNA improves the accuracy of predicted expression.

We next scrutinized the features used by our models to identify the specific sequences that define cell-type specific gene expression. As expected, many motifs that predict expression were also detected as enriched in accessible chromatin (**Fig. 1G**). Examples include motifs for SPIC and SPI1 in the macrophage model, MEF2A in the cardiomyocyte model, SOX9 in the vascular endothelial and endocardium models, and

CEBPA in the fibroblast model (**Fig. 5D**). The models also explicitly identified motifs that are predictive of reduced expression, identifying putative respressors. For example, motifs for HEY2 - which contributes to cardiomyocyte specification in development ^78^ - were inversely associated with expression in cardiomyocytes and other cell types (**Fig. 4D**). This result agrees with the factor’s known role as a transcriptional repressor ^79^, and would be consistent with HEY2 playing a role in other cell types apart from its characterized function in cardiomyocytes ^78^. Similarly, the models for T cells and mast cells all captured an inverse relationship between expression and SMAD4 (**Fig. 5E**). TGFB signaling via nuclear translocation of SMAD4 is highly cell-type specific, driving a broadly immune-suppressive role ^80^ with SMAD4 alternately acting as a transcriptional repressor or activator depending on cellular context ^81^. In addition, we observed cases where a particular TF motif was utilized by models across cell types, such as ZNF384 and YY1 motifs leading to increased expression predictions in all cell types (**Fig. 5D**). Such relationships are difficult if not impossible to detect in a testing strategy looking for enrichments in a cell type compared to others. In summary, predictive models identify TFs that play cell type-specific roles, TFs related to expression across many cell types, and assign an explicit direction of effect on transcription.

## Discussion

We generated a resource of snATAC- and snRNA-Seq from multiple donors and utilized a state-of-the-art regression approach to find alterations by sex and age. We additionally studied regulatory roles of transcription factors in distinct cell populations through enrichment analyses and predictive expression models. Performing single-cell or single-nucleus RNA analysis in solid human tissue is difficult, and our dataset represents one of only a small handful of studies covering the healthy human heart at single-cell resolution from multiple donors ^6–8,17^. Our data allows us to characterize alterations at multiple levels, such as finding sex-specific TGFB-driven transcription (**Fig. 2B**) accompanied by altered epigenetic signatures of TGFB effectors and regulators (**Fig. 2F**). Cell-type specific changes - such as statistically significant TNFA activation hallmarks in male T cells but not other cell types (**Fig. 2B**) - and altered cell type abundances (**Fig. 2E and 3E**) highlight the key role of single-cell resolution for understanding alterations by donor traits. Such cell type-specific changes could contribute to why a recent analysis of sex-dependent bulk transcriptional changes found most sex-specific changes to be of small magnitude and detected a minimal number of statistically significant pathway alterations in heart samples ^82^. Our results indicate that single cell libraries from a tractable number of donors can identify diverse alterations in cell type proportions, transcriptional differences, and epigenetic programs varying by donor traits.

In addition to facilitating tests of sex- and age-dependent differences across cell types, our dataset also allowed us to quantify the level of transcriptional variation that can be accounted for using a simple, binary TF motif-based linear model when using or excluding TSS-distal sequence (**Fig. 5B**). Our results underscore the importance of distal regulatory information in determining gene expression levels ^74,77,83^ and reaffirm the observation that - at least in a simple modeling framework - simple models using TF motifs as input features explains only a minority of total variation in RNA levels ^72,73^. Furthermore, our models identify regulatory factors through a method that complements widely-used tests ^8,17,23^ based on motif presence in accessible peaks (**Fig. 1F**). In total, our predictive models of expression extend existing analyses of cell-type specific regulatory programs in the adult human heart ^8,17^ and establish a baseline level of accuracy for comparison against more complex methods ^20–22^ utilizing larger datasets and less interpretable models.

Our analyses come with several caveats, most notably due to the limited number and diversity of samples analyzed. While we find statistically significant alterations by age and sex within the samples of our dataset, the limited number of individuals studied precludes generalizing those differences to the population at large. Considerably larger datasets will be necessary to find generalizable differences while accounting for confounding variables like donor medical history and disease status, dissect finer-grained effects like sex-specific differences before or after menopause onset ^84^, and provide statistical power to detect biologically meaningful changes of small magnitude. In addition, analyses of new datasets - or meta-analysis of existing ones - will be required to study differences occurring in cell populations like atrial cardiomyocytes ^6,7^ that we did not study because of the limited anatomical coverage of our samples. In our predictive models of expression, due to limited training inputs and a preference for easily interpretable models we employed a simple linear model based on motif presence/absence. Future work in more complex - and likely more accurate - models like deep neural networks ^20–22^ will be required to understand the relative value of distal DNA sequence and of particular sequence features/motifs in alternate prediction frameworks. It will be particularly interesting to test the reproducibility of cell-type-specific TF roles found via our linear models (**Fig. 5D**), given inherent co-occurence of motifs in the genome (**Supplemental Fig. 5D**). As modeling approaches that use regularization to select a minimal set of predictors (e.g. LASSO and elastic net regression) will often pick only one member of a motif “family”, motif/expression relationships in particular cell types will require validation in further modeling approaches.

Our findings raise a number of natural avenues for further research. First, analyses of larger datasets of single-cell heart tissue - either newly generated, or via meta-analyses of existing datasets - will be crucial for identifying sex- and age-dependent variation that generalizes to the population at large, and to extend analysis to cover other important patient covariates or interactions. Relatedly, similar analyses in other organs can study the extent to which sex- and age-dependent variation observed in cell types of the heart is reproduced in cell types of other human tissues. Furthermore, perturbational work in animal or culture models will play a crucial role in studying differences detected in large-scale, single-cell analyses by avoiding limitations inherent to observational designs. Our identification of biological processes such as metabolic shifts, TGFβ signaling, and inflammation also raise the prospect that larger analyses may detect alterations by sex or age that are already of clinical interest in cardiac disease ^85–87^ and raise the possibility of personalizing therapies by patient traits. Reassuringly, our identification of numerous statistically significant differences in both RNA- and ATAC-seq data using a modest number of donors bodes well for the ability of future analyses to identify molecular covariates of patient traits within datasets that can feasibly be generated with current technology.

## Methods

### Tissue Collection

This study complies with all relevant ethical regulations and was approved by the University of Washington Institutional Review Board (STUDY00002144). Informed consent was obtained prior to collection of human tissues. No compensation was provided for participation. Collected samples were absent of evidence of disease upon review by study clinicians. Details regarding the collection are available on protocols.io ^88,89^.

### Single-nucleus Library Generation

Nuclei for sci RNA-Seq were extracted from frozen, powdered heart tissue. 200-250 mg of frozen tissue was powdered while frozen, then dissociated using a Gentle MACS Tissue Dissociator at 4C using 5 mL of ice-cold lysis/fixation buffer containing 10 mM sodium phosphate (pH 7.2), 3 mM MgCl^2^, 10 mM NaCl, .02% Triton X-100, 5% glutaraldehyde, 1% DEPC, 10 mM ribonucleoside vanadyl complex (NEB). Dissociated tissue was filtered through a 70 uM cell strainer on ice and washed with an additional 5 mL of ice-cold lysis/fixation buffer. The buffer/nuclei mixture was then incubated for 15 minutes at 4C in a rotating 15 mL Falcon Tube. Nuclei were pelleted by centrifugation at 600 RCF for 8 minutes at 4C. Supernatant was decanted, then nuclei were resuspended in 1 mL of nuclei suspension buffer (NSB) containing 10 mM Tris HCl, pH 7.4. 10 mM NaCl, 3 mM MgC l^2^, 1% SuperaseIn, 1% bovine serum albumin (BSA) solution (NEB, 20 mg/mL). Nuclei were pelleted at 600 RCF for 5 minutes at 4C, and supernatant was decanted. Nuclei were resuspended in 100 uL of NSB per aliquot, then snap-frozen with liquid nitrogen.

Libraries were generated using a 3-level sci RNA-Seq workflow ^24^. The workflow was modified to add a FACS sorting step following ligation in order to minimize background RNA levels, with a detailed workflow available at protocols.io (https://www.protocols.io/view/3-level-sci-rna-seq-with-facs-dm6gpw255lzp/v1). Libraries were sequenced using an Illumina Nextseq 500 high output sequencing kit.

Nuclei for sci ATAC-Seq were extracted from powdered, frozen tissue and fixed as in previous work ^23^. Libraries were generated using a 3-level sci ATAC-Seq workflow ^23^ and sequenced using an Illumina Novaseq sequencer.

### Single-nucleus RNA-Seq Analysis

Raw sequencing output was processed using a pair of Nextflow processing pipelines available at https://github.com/bbi-lab/bbi-dmux (handling sample demultiplexing) and https://github.com/bbi-lab/bbi-sci (handling assignments of reads to cells, filtering, alignment, and cell-by-gene matrix generation).

Analysis of single-cell RNA-Seq data was performed using Monocle 3 ^24^. Cells were filtered by discarding any with unique molecular identifiers (UMIs) less than 100, mitochondrial RNA percentage greater than 10, or a Scrublet doublet score ^90^ of over .2. A 2-dimensional UMAP representation ^91^ of cells was found after using mutual nearest neighbors alignment ^92^ to align by sample. Cell type assignments were made manually based on expression of marker genes in UMAP clusters (**Figure 1D**). Plots of several QC metrics - overlaid over 2D UMAP embeddings or shown as distributions per sample - are available in **Supplementary Fig. 1**.

### Single-nucleus ATAC-Seq Analysis

Sequencing output was processed using a pair of Nextflow processing pipelines available at https://github.com/bbi-lab/bbi-sciatac-demux (handling demultiplexing) and https://github.com/bbi-lab/bbi-sciatac-analyze (assigning reads to cells, aligning reads, calculating peaks, finding motif occurrences in peaks, and generating cell x peak matrices).

Analysis of single-cell ATAC-Seq data was performed using Monocle 3 ^24^. Cells were filtered by discarding any with unique molecular identifiers (UMIs) less than 1000, fractions of reads in TSS (FRIT) of less than .08, fractions of reads in peaks (FRIP) less than .2, or a doublet likelihood of greater than .5 ^90^. Gene activity scores were calculated using ArchR ^93^ using default settings. Cell-by-gene activity score matrices were then used to generate a Monocle CDS object. The ATAC-Seq data was then aligned with the filtered RNA-Seq data using Harmony ^94^ based on all genes shared between RNA and ATAC CDS objects, and a new UMAP embedding was generated based on the corrected PCA coordinates of both datasets after Harmony correction. Based on UMAP coordinates in this new embedding, ATAC-Seq cells were labeled using a k-nearest neighbor transfer from the k=7 nearest RNA-Seq cells (using cell assignments described above for RNA-Seq data). Plots of several QC metrics - overlaid over 2D UMAP embeddings or shown as distributions per sample - are available in **Supplementary Fig. 2**.

### Differential motif abundance testing in accessible peaks

The presence of TF motifs in peaks was calculated based on the presence of any motif occurrence in the peak DNA sequence below a p-value cutoff of 1e-7 using MOODS ^95^. Motif count x cell matrices were then made by multiplying a motif (rows) x peaks (columns) matrix with a peak (rows) x cell (columns) matrix, generating a motif-count x cell matrix where each entry corresponded to the number of peaks accessible in a given cell that contained a given TF motif.

To test for motif abundances that varied by a function of donor covariates (age and sex), testing was run separately for all cells of a single cell type. Testing for motif counts was done using a GLMM fit using the lme4 package ^96^, using a negative binomial model with sample donor as a random effect, as well as fixed effects of anatomical site, donor age, and sex. Multiple testing correction was performed with the Benjamini-Hochberg procedure ^97^. This modeling approach is available in current releases of Monocle 3 ^24^.

For testing of cell type-specific motif enrichments (**Fig. 1F**), testing was run for all cells at once. To test for motifs enriched in a specific cell type, all cells were assigned a dummy variable valued as ‘1’ for cells that are from the type being tested, and ‘0’ for all others. Testing was then run using a GLMM fit using the lme4 package ^96^, using a negative binomial model with sample donor as a random effect, as well as fixed effects of the cell-type-dummy variable, anatomical site, donor age, and sex. Multiple testing correction was performed with the Benjamini-Hochberg procedure ^97^.

### Differential expression testing

DE testing used a GLMM fit using the lme4 package ^96^, using a negative binomial model with sample donor as a random effect, as well as fixed effects of anatomical site, donor age, and donor sex. This modeling approach is available in current releases of Monocle 3 ^24^. Multiple testing correction was performed with the Benjamini-Hochberg procedure ^97^.

Gene set enrichment analysis tested for enrichments by age or sex within 50 Hallmark Pathways accessed from the MSigDB collection ^98^ accessed through the msigdbr R package. Testing used the fgsea package ^99^ and multiple testing correction was performed with the Benjamini-Hochberg procedure ^97^.

### Adult versus fetal enrichment comparisons

Enrichments for TF motifs in accessible chromatin of fetal cell types was accessed at https://descartes.brotmanbaty.org/bbi/human-chromatin-during-development/ (see “Motif enrichment across cell types” section for download link). Enrichments in adult cell types were calculated as described above under “Differential motif abundance testing in accessible peaks”. Comparisons were made between the following adult-to-fetal matchings: “Cardiomyocyte” and “Cardiomyocytes”; “Vascular Endothelium” and “Vascular endothelial cells”; “Endocardium” and “Endocardial cells”; “Macrophage” and “Myeloid cells”; “Perivascular Cells” and “Smooth muscle cells”; “Fibroblasts” and “Stromal cells”; “Adipocytes” and “Epicardial fat cells”; “Neuronal” and “Purkinje neurons”; “T Cells” and “Thymocytes”.

For each comparison, plots (**Fig. 4A-D; Supplementary Fig. 4; Supplementary Table 7**) were calculated using only motifs that were significantly enriched in either adult or fetal data at a FDR cutoff of .1. Outliers were selected based on qualitative divergence from broad cell type correlations in enrichments between fetal and adult cells.

### RNA Expression Predictive Modeling

First, pseudo-bulk expression levels were calculated by pooling all UMIs for all genes for cells within a particular cell type. These were used to quantify the transcripts per million for each gene. Log2(TPM) was then used as the RNA expression level to be predicted for a particular gene/cell type pair.

To link distal sites to promoters, we ran Cicero ^77^ to quantify covariance among peaks across all cell types. To link distal sites to genes, we first defined any peaks that intersected a defined window around the TSS (this region size was a hyperparameter set through performance on a validation set, see below). Then, any peaks outside the promoter set of peaks that were linked with a co-accessibility score greater than some cutoff (a hyperparameter). Motifs from the JASPAR database ^100^ “2018 Non-redundant Vertebrates” motif set were determined using FIMO ^101^ at varying p-value cutoffs. For models using promoter sequence only, features would be a binary value for if one or more motif occurrences was found in the promoter sequence below a p value cutoff. For models using promoter and distal sequence, features were binary values for if a motif occurred in the promoter or distal regions.

RNA expression for protein-coding genes was predicted with an elastic net linear model using motif presence/absence as features. Data was divided into train, test, and validation sets at the level of chromosomes (approximately a 80/10/10 split in gene numbers) for hyperparameter setting. First, promoter size and motif p values were varied (Promoter sizes upstream/downstream of TSS were 1000/200, 1500/500, and 5000/2000. P values tested were 1e-4, 1e-5, 1e-6). Models were trained on the training set setting l1/l2 penalties by internal cross-validation, then evaluated on the validation set. The best average performance occurred using a p value cutoff of 1e-4 with a promoter of 1500/500 bases upstream/downstream. Holding that promoter region size constant, we trained models varying the cicero co-accessibility cutoff to link a distal site, maximum number of distal sites to link, and window size of DNA bases to scan centered at a linked peak. We tested combinations of: Co-accessibility cutoffs of .015, .035, .05; max distal sites of 5, 10, or 20; distal sit size of 600 or 1,000 bases, motif p value cutoffs of 1e-4, 1e-5, and 1e-6. Models were trained on the training set setting l1/l2 penalties by internal cross-validation, then evaluated on the validation set (**Supplementary Fig. 5A**). Optimal performance was obtained using a co-accessibility cutoff of .015, a maximum of 5 distal sites, 1000 bp distal site windows, and a motif cutoff of 1e-4. Those parameters were then set for use in training a model for evaluation on the test set. L1/l2 penalties were set by internal cross-validation on a pooled training + validation set, then a model was trained using those penalties and the best hyperparameters found earlier. The model was then evaluated on the test set (**Fig. 5B**). Finally, holding those hyperparameters constant, a final model was trained using all three train, validation, and test sets. The coefficients of this final fit model are reported in **Fig 5D**.

## Code Availability

Code is accessible in a Github repository that will be made public upon publication. A copy of the repository at the time of submission has been made available to reviewers and is available upon request.

## Data Availability

Data is available through the HuBMAP consortium website’s data portal (https://portal.hubmapconsortium.org/). Dataset IDs for samples presented in this study are available in **Supplementary Table 8**.

## Acknowledgments

We thank the tissue donors and their families, without whom this work would be impossible.

We thank Andrew Hill and Silvia Domcke for their contributions to single cell ATAC-Seq processing pipelines and computational methods. We thank Haleigh Larson and Christian Pfleger for their work developing nuclei isolation and fixation methods. We also thank the team at the Northwest Genomics Center for their contributions to library sequencing.

Funding for this work was provided by NIH U54 grant U54HL145611, CZI HCA Seed Networks CZF2019-002442 and Inflammation CZF2020-218446 awards, and the Brotman Baty Institute for Precision Medicine. JS is an Investigator of the Howard Hughes Medical Institute.

## Author Contributions

D.F.R., G.B., R.D., D.J., R.G., S.S., A.R.G., and D.O. performed sample processing and library preparations; Y.L. collected samples; D.F.R., G.B., and R.D. ran analysis; D.F.R., B.E., J.F., C.S., A.A.G., and B.M. developed methods, software, and protocols; L.S., J.S., S.L., C.T. supervised work Beale, A. L., Meyer, P., Marwick, T. H., Lam, C. S. P. & Kaye, D. M. Sex Differences in Cardiovascular Pathophysiology: Why Women Are Overrepresented in Heart Failure With Preserved Ejection Fraction. *Circulation* **138**, 198–205 (2018).

