## Supplemental_Figures for "Single-cell analysis of chromatin and expression reveals age- and sex-associated alterations in the human heart"

Supplementary Figure 1

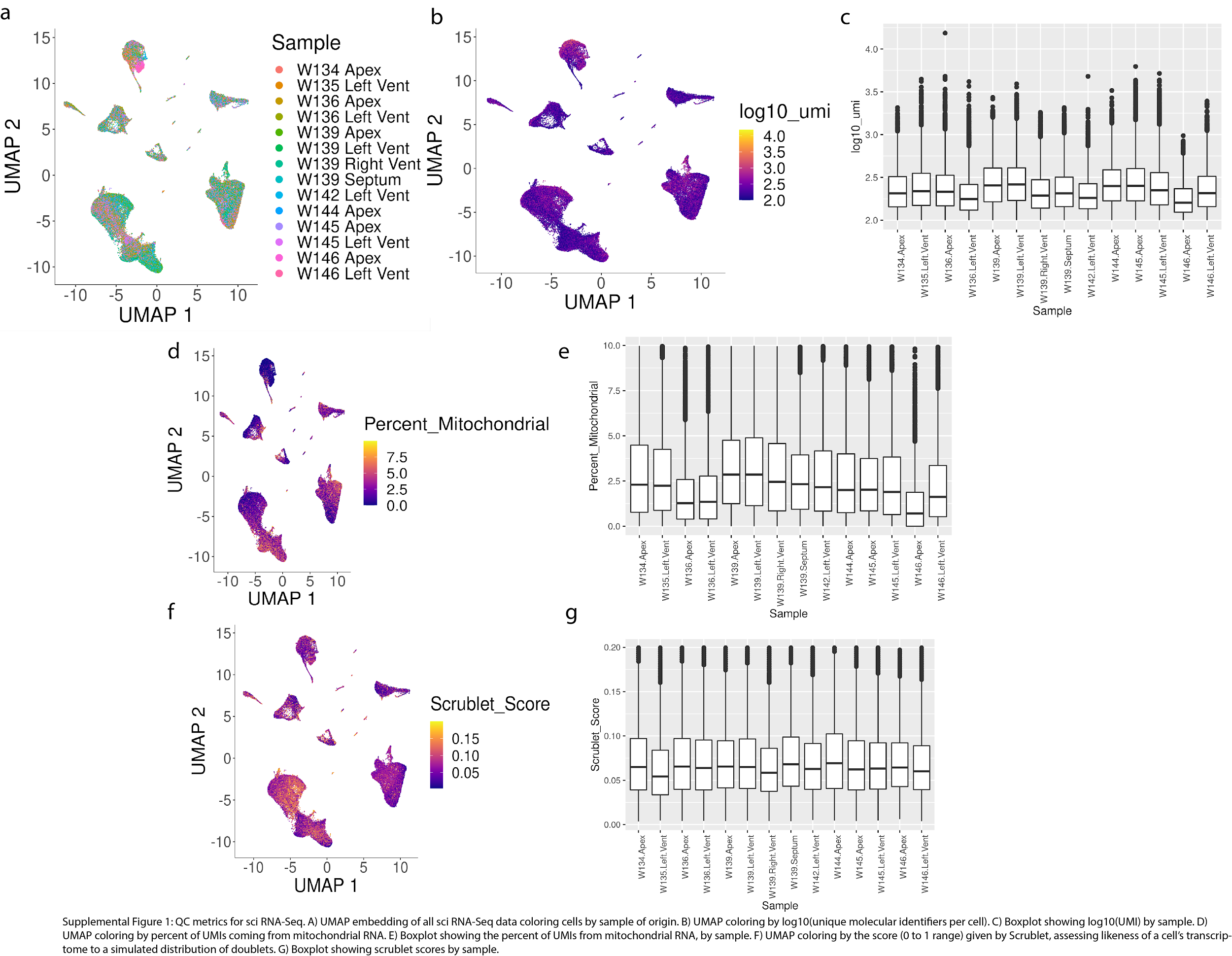

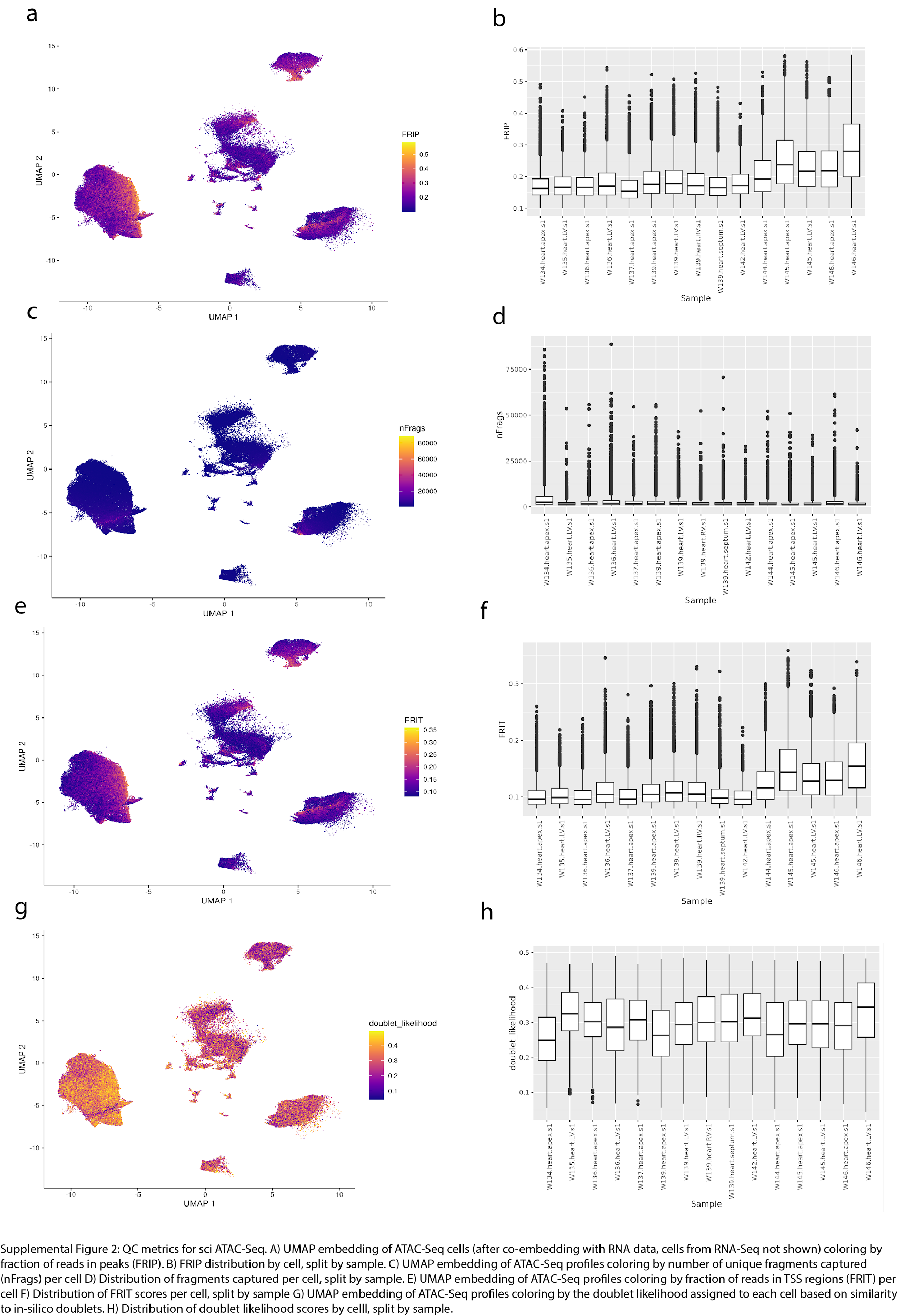
Supplementary Figure 2

Supplementary Figure 3:

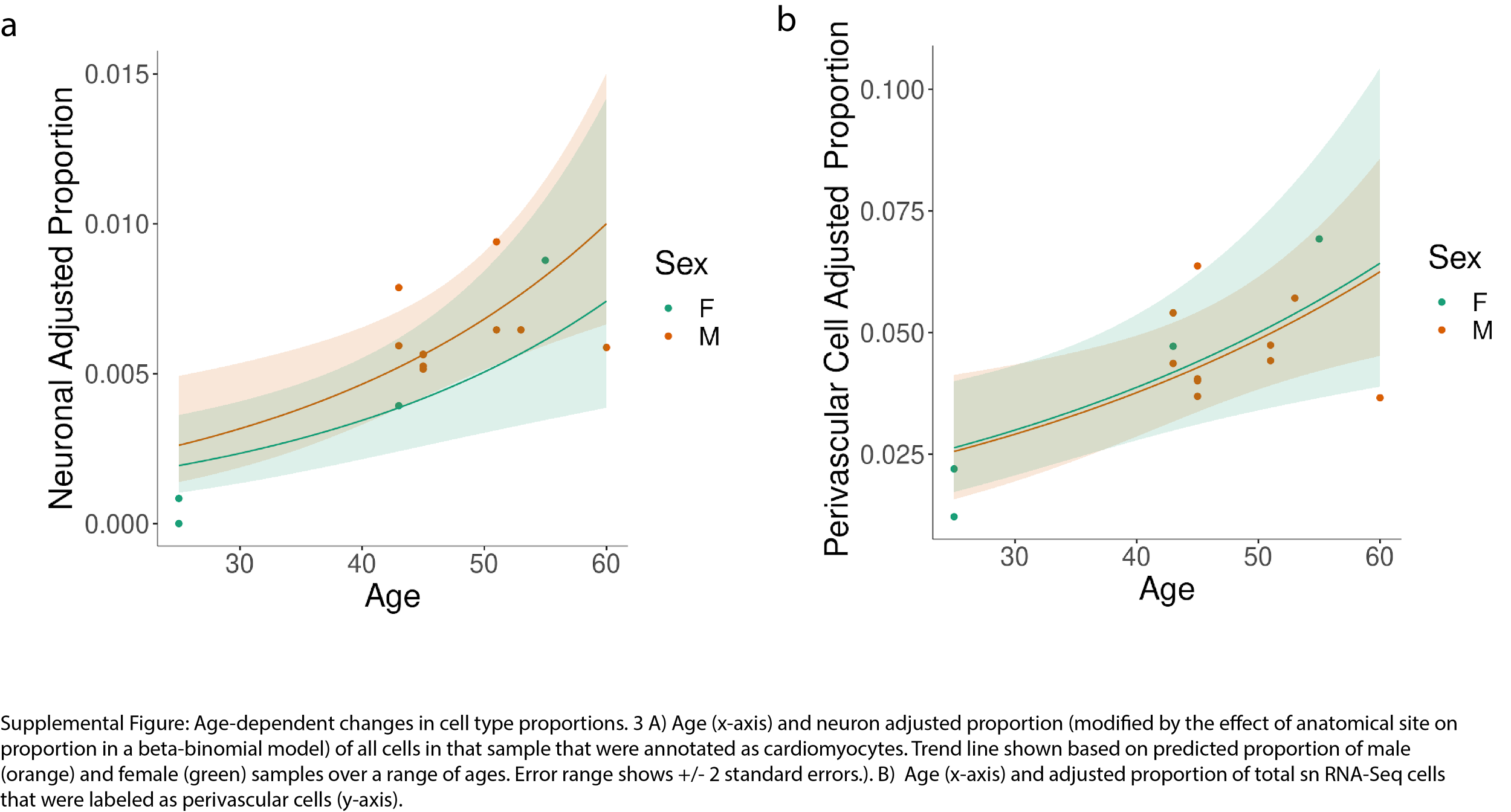

Supplementary Figure 4:

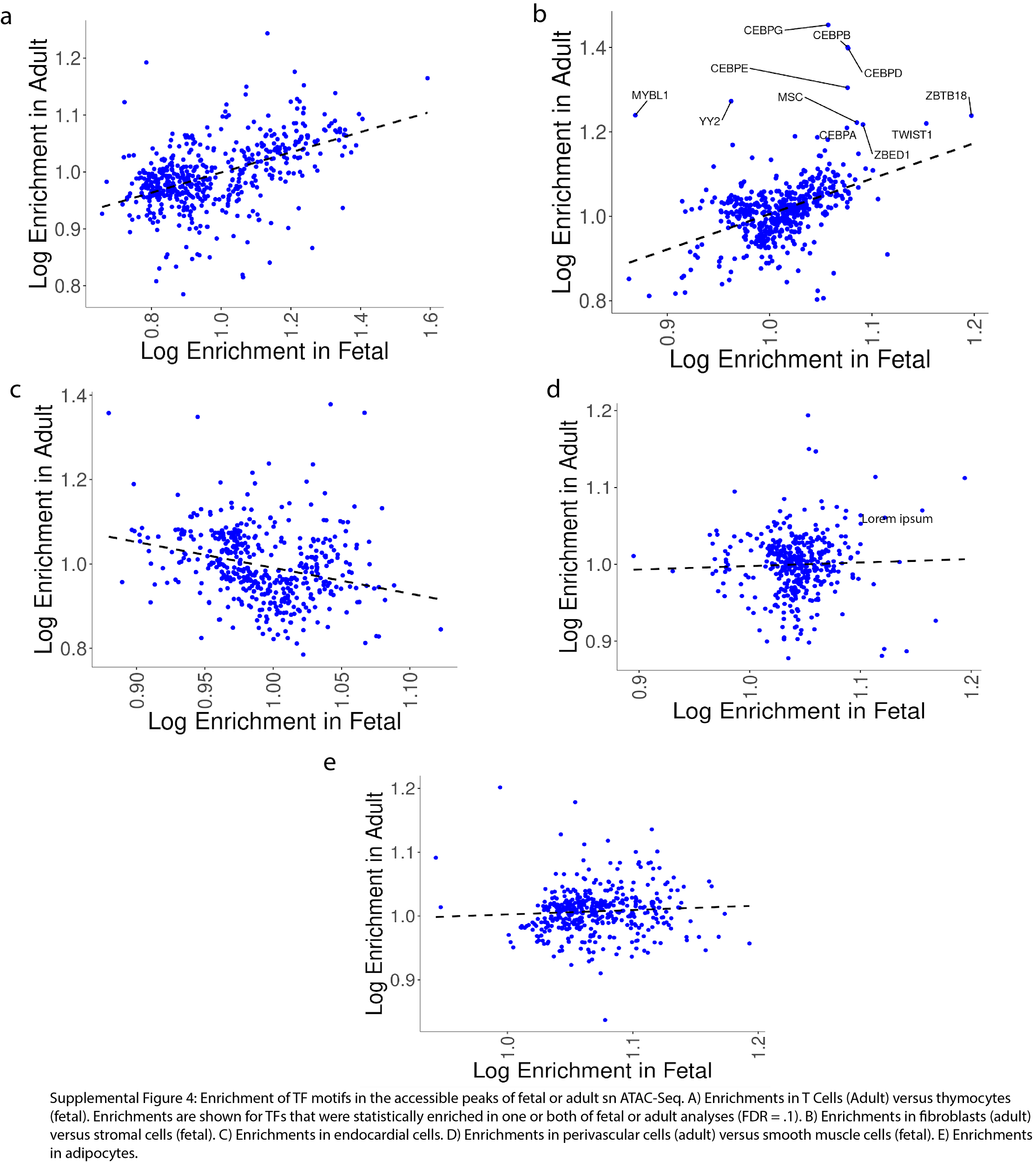

Supplementary Figure 5:

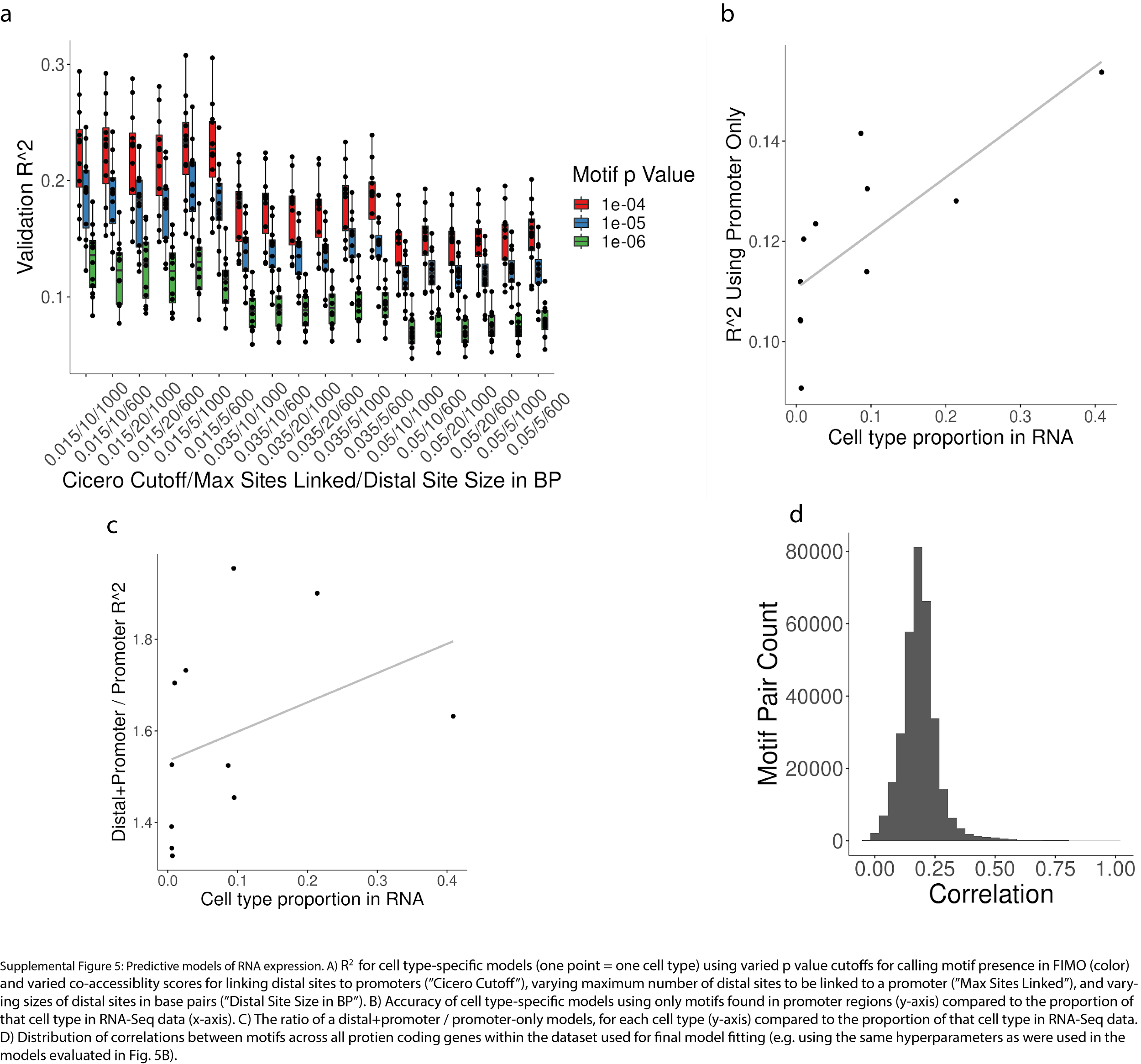
